# The membrane-actin linker ezrin acts as a sliding anchor

**DOI:** 10.1101/2021.11.28.470228

**Authors:** Elgin Korkmazhan, Alexander R. Dunn

## Abstract

Protein linkages to filamentous (F)-actin provide the cell membrane with mechanical resiliency and give rise to intricate membrane architectures. However, the actin cytoskeleton is highly dynamic, and undergoes rapid changes in shape during cell motility and other processes. The molecular mechanisms that underlie the mechanically robust yet fluid connection between the membrane and actin cytoskeleton remain poorly understood. Here, we used a single-molecule optical trap assay to examine how the prototypical membrane-actin linker ezrin acts to anchor F-actin to the cell membrane. Remarkably, we find that ezrin forms a complex that slides along F-actin over micron distances while resisting mechanical detachment. The ubiquity of ezrin and analogous proteins suggests that sliding anchors such as ezrin may constitute an important but overlooked element in the construction of the actin cytoskeleton.

## MAIN

Protein linkages between the cell membrane and the actin cytoskeleton provide the cell membrane with mechanical resiliency (*1*–*3*) and give rise to intricate membrane architectures such as microvilli in the intestinal epithelia (*3*–*6*). However, the actin cytoskeleton is also dynamic on the seconds timescale, a property that underlies its ability to drive membrane shape changes (*1*) during phagocytosis (*7, 8*) and amoeboid motility (*9*). Even apparently static structures such as microvilli are maintained via constant F-actin treadmilling (*10*). At present, it is unclear how a mechanically robust yet fluid connection between the membrane and actin cytoskeleton is maintained in these diverse circumstances (*3, 11*).

A general assumption is that numerous yet weak, transient crosslinks between filamentous actin (F-actin) and the membrane are responsible for this phenomenon (*3, 12, 13*). An unexamined, alternative possibility is that proteins linking the membrane and actin cytoskeleton might respond differently to forces oriented parallel vs. perpendicular to the membrane plane, potentially allowing F-actin to slide relative to the membrane while maintaining a mechanically stable attachment. Neither of these possibilities have been subject to a direct experimental test. More broadly, to our knowledge no study to date has systematically examined the response of F-actin binding proteins to load oriented parallel vs. perpendicular to the F-actin filament, a distinction that might be expected to be critical in the case of cytoskeletal membrane anchors.

In this study we focused on ezrin as a prototypical membrane to F-actin crosslinker. The ezrin-radixin-moesin (ERM) protein family (*4, 14*) emerged prior to the divergence of choanoflagellates and metazoans, and its members are present in all sequenced animals (*15*). ERMs link the cell membrane to the actin cortex (Fig. 1a), a thin meshwork of F-actin and myosin II that gives the cell membrane mechanical resiliency (*1, 4, 12*). In addition, ERMs stabilize membrane protrusions such as the microvilli in the gut (*6, 10, 16*–*18*) and retina (*19*), and play key roles in regulating cell shape change (*20*–*23*) and signal transduction (*4, 5, 17, 24*–*28*). All three ERM proteins are likely to experience mechanical load as part of their physiological functions. For example, ezrin helps drive compaction in the early mammalian embryo (*6, 29*–*31*), reinforces membrane to F-actin attachment during bleb retraction (*2*), and localizes to microvilli where membrane-actin forces are anticipated to be high (*10*). Consistent with these functions, ezrin knockout mice are born at submendelian ratios and do not survive past weaning due to a failure to form functionally adequate intestinal microvilli (*6*). How ezrin maintains a mechanically stable yet dynamic linkage to F-actin is, to our knowledge, not understood.

**Figure 1.**
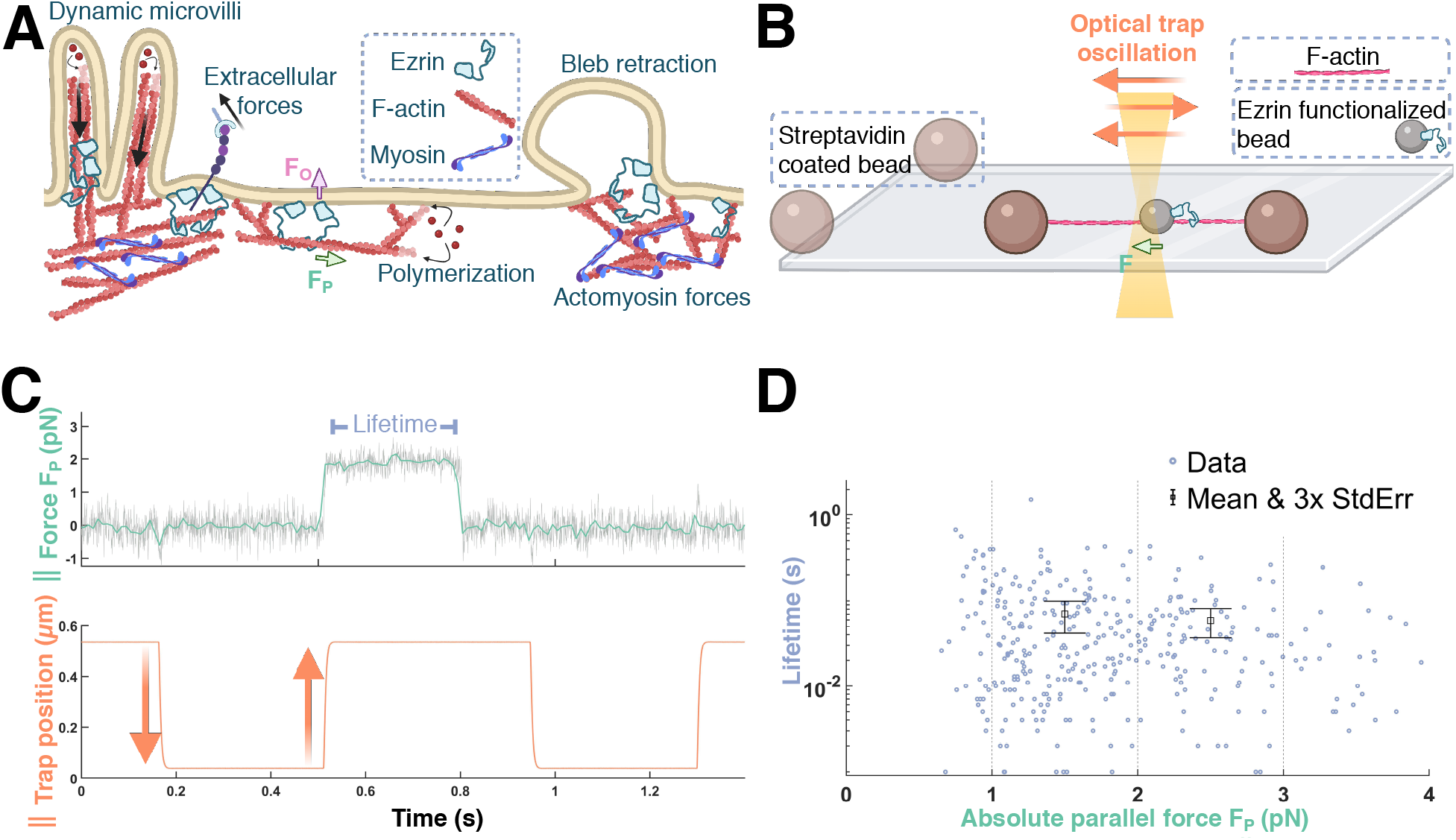
Tightrope assay reveals the force-dependent interaction between F-actin and single ezrin-T567D molecules. **(a)** Ezrin links the cell membrane to the actin cortex, anchors the bases of microvilli, and aids in membrane bleb retraction. In each case, ezrin must resist force orthogonal to the membrane (*F*_*O*_) while allowing F-actin flow in response to force parallel to the membrane surface (*F*_*P*_). **(b)** Cartoon of the tightrope assay (not to scale). Single actin filaments are tautly suspended between surface-attached beads. A bead functionalized with ezrin-T567D (here, a single molecule) is captured with the optical trap and oscillated along or perpendicular to the filament. A binding event forces the bead out of the trap center, after which the oscillation is immediately paused to measure the bond lifetime at the given force. The oscillation is resumed following unbinding. **(c)** Example force trace at 100 Hz (*green*) and 4 kHz (*gray*), for binding of a single ezrin-T567D molecule to F-actin under load parallel (∥) to the filament axis. **(d)** Bond lifetimes upon parallel loading of single ezrin-T567D molecules (352 events, from 8 beads from 8 flow cells). The mean lifetime is <100 ms for 1-3 pN forces. Mean and error bars (3 standard errors of mean) for 1-2 and 2-3 pN are plotted.

## RESULTS

As with other ERM proteins, ezrin attaches to the membrane by binding to the membrane lipid phosphatidylinositol 3,4-bisphosphate (PI(4,5)P_2_) and/or to protein partners. A key step in ezrin activation is the phosphorylation of a conserved threonine residue (T567 in human ezrin), which is believed to help free its C-terminal F-actin-binding region (*4, 32, 33*). The phosphomimetic mutant (ezrin-T567D) has been used extensively to study activated ezrin *in vivo* and *in vitro*. We thus used human ezrin-T567D as a model with which to determine how the interaction of ezrin with F-actin responded to the magnitude and orientation of applied mechanical load.

We adapted a single-molecule optical trap assay (*34*), hereafter termed the “tightrope” assay (*35*), that allows us to apply force to ezrin-T567D bound to filamentous (F)-actin both parallel and perpendicular to the filament axis (Fig. 1b; Methods). In this assay, streptavidin-coated, 3 μm-diameter beads are adhered to the surface of the coverslip. Biotinylated fluorescent actin filaments are tautly suspended between pairs of beads via the flow generated by rapidly pipetting the F-actin solution through the flow cell. After excess F-actin is washed out, 1 μm beads sparsely functionalized with ezrin-T567D fused to a HaloTag domain (6xHis-HaloTag-ezrinT567D) are added to the flow cell, along with ∼2 μM of a PI(4,5)P_2_ analog (Methods), whose presence helps to activate F-actin binding. An ezrin-functionalized bead is captured in an optical trap and is then oscillated along or perpendicular to an actin filament. A binding event results in force that pulls the bead out of the optical trap center, which we measure with ∼1 ms and ∼0.1 pN precisions (Fig. 1c). Upon detection of a binding event the trap oscillation is temporarily stopped and the lifetime of the bond at the bound force is recorded (Methods).

We first examined the effect of parallel load on the interaction of single molecules of ezrin-T567D with F-actin. To ensure that a given optically trapped bead that showed binding activity most likely contained only one active molecule, we labeled beads with low concentrations of ezrin-T567D, such that ∼90% of beads showed no binding activity (Methods). Consistent with Poisson statistics (Methods; Supp. Table 1) experiments at this labeling ratio revealed that ∼8% of all tested beads exhibited either solely single step unbinding events or a mixture of single- and double-step unbinding events, as expected from beads containing 1 and 2 ezrin-T567D molecules respectively, where each molecule had a single actin binding site (Fig. 1c). We thus interpreted the events from beads exhibiting solely single step unbinding to be from a single ezrin-T567D molecule interacting with the actin filament. Measurements from such beads revealed a particularly weak interaction, with a <100 ms mean binding lifetime when bearing 0.5-4 pN, forces comparable to those generated by individual myosin motor domains (*36*–*38*) (Fig. 1d). These lifetimes are consistent with atomic force microscopy experiments that inferred the mean actin binding lifetime at zero load to be <1 second (*13*).

*In vivo*, ezrin is found as oligomers, and more generally in clusters, in addition to monomeric forms (*39*–*41*). Thus, we next tested the binding of multiple ezrin-T567D molecules to F-actin. We increased the labeling ratio of beads such that ∼35% of beads showed F-actin binding activity (Methods). According to Poisson statistics, at this labeling ratio ∼80% of the active beads are expected to be labeled with a single ezrin-T567D molecule, ∼17% are expected to contain 2 molecules, and 3% more than 2. Consistently, we mostly observed single step and double-step unbinding for active beads, as expected from beads binding to F-actin with one or two molecules at a time (Supp. Table 1; Methods). However, ∼14% of active beads (18 out of all 404 beads tested), showed a qualitatively different behavior, in which a step in applied load relaxed back to 0 - 0.1 pN while the complex was still bound, as opposed to exhibiting step-unbinding (Fig. 2a). This behavior occurred repeatedly for a given bead.

**Figure 2.**
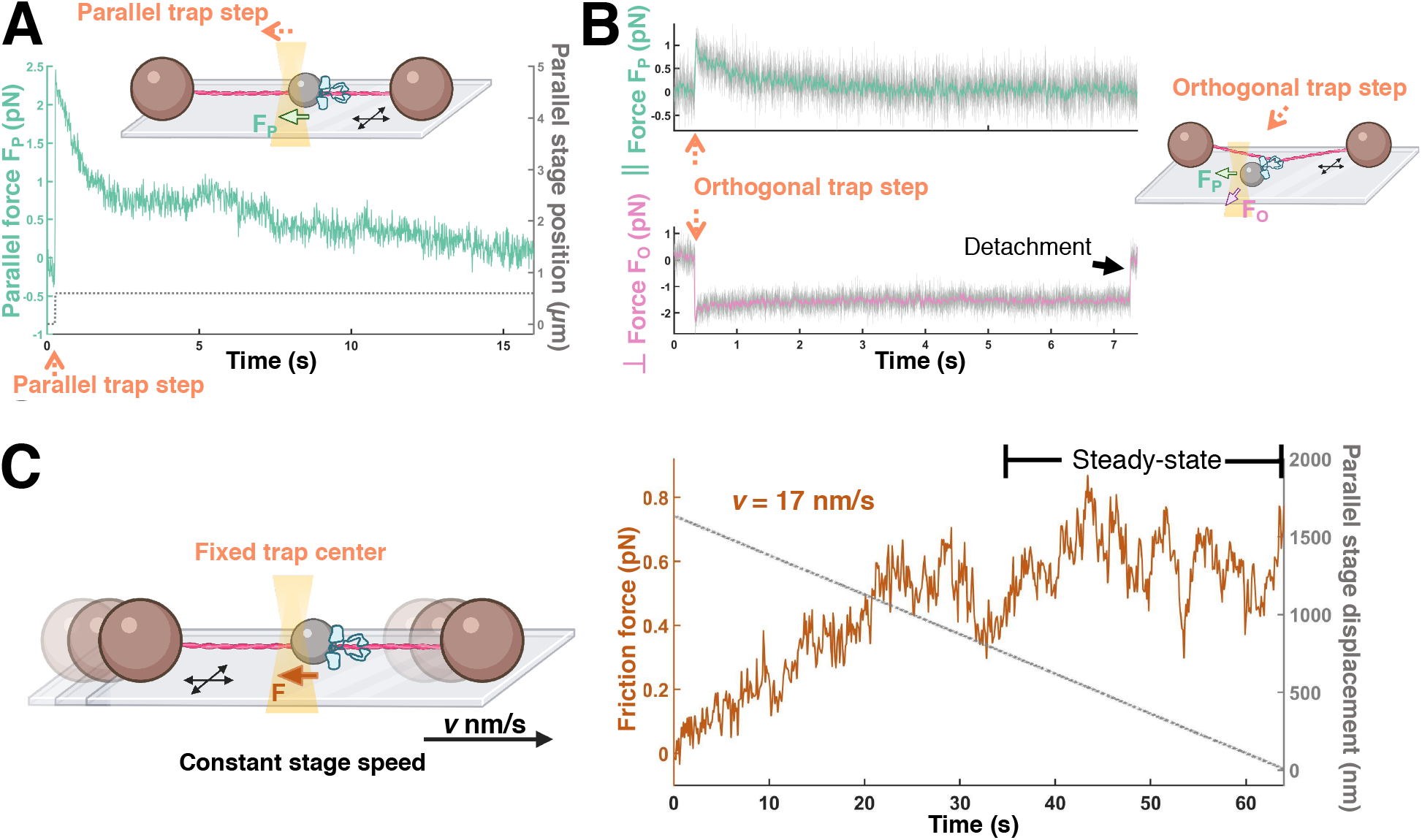
Multiple ezrin-T567D molecules form a sliding anchor on F-actin. **(a)** Force relaxation from ∼2 to 0.1 pN following a step in parallel (∥) load, and **(b)** from ∼1 to 0.1 pN in the presence of a simultaneous orthogonal (⊥) load. Complexes slide along F-actin before unbinding, seen as a step in the orthogonal force trace. Parallel and perpendicular force traces are at 100 Hz (*green* and *magenta* respectively) and 4 kHz (*gray*; shown only in (b)). **(c)** Ezrin-T567D complexes slide for multiple microns along F-actin. *Left*: the optical trap is held in a fixed position while the microscope stage moves at a fixed velocity. *Right*: this results in a frictional force on the ezrin-T567D complex that asymptotes to a steady-state. Force trace is at 10 Hz.

The statistical rarity of beads exhibiting this force relaxation behavior suggested that it arose from multiple ezrin-T567D molecules acting in concert (Methods). Importantly, such complexes could still relax force parallel to the filament axis to ∼0 pN even in the presence of perpendicular load (Fig. 2b). This observation suggested the ability of the complexes to slide along the filament without transient detachment events that would otherwise result in irreversible detachment in the presence of perpendicular load. Consistently and remarkably, we found that these complexes remained stably associated with F-actin for tens of seconds and over micron distances when the stage was translated at constant velocity, and that they could repeat this behavior for multiple such ramps in succession (Supp. Fig. 1) before exhibiting step-unbinding from the filament.

To determine the characteristics of the minimal ezrin assembly that would support sliding, we next performed experiments at limiting ezrin-T567D labeling concentrations, such that sliding was rare (6 of 289 beads examined). Of these, five beads, from independent bead preparations, produced matching steady-state friction forces for a given velocity (Fig. 3a; Methods, Supp. Fig. 2) and exhibited step-unbinding. Both observations are consistent with the presence of a defined, minimal ezrin-T567D cluster with set composition and behavior. Similar steady-state friction values and single-step detachment were likewise observed for a subset of beads labeled with higher ezrin-T567D concentrations (60% of all sliding beads, 2% of all beads). Data pooled from these complexes yielded narrowly distributed steady-state friction values that increased monotonically with increasing velocity, as would be expected from a well-defined complex (Fig. 3b). The binding lifetimes for the minimal sliding complex placed under orthogonal load were well-described as a slip bond with an extrapolated mean lifetime at zero load of ∼80 s (Fig. 3c), again consistent with the presence of an underlying homogeneous population. In sum, these observations support the presence of a minimal complex that is homogenous in nature and able to slide along F-actin for tens of seconds without unbinding.

**Figure 3.**
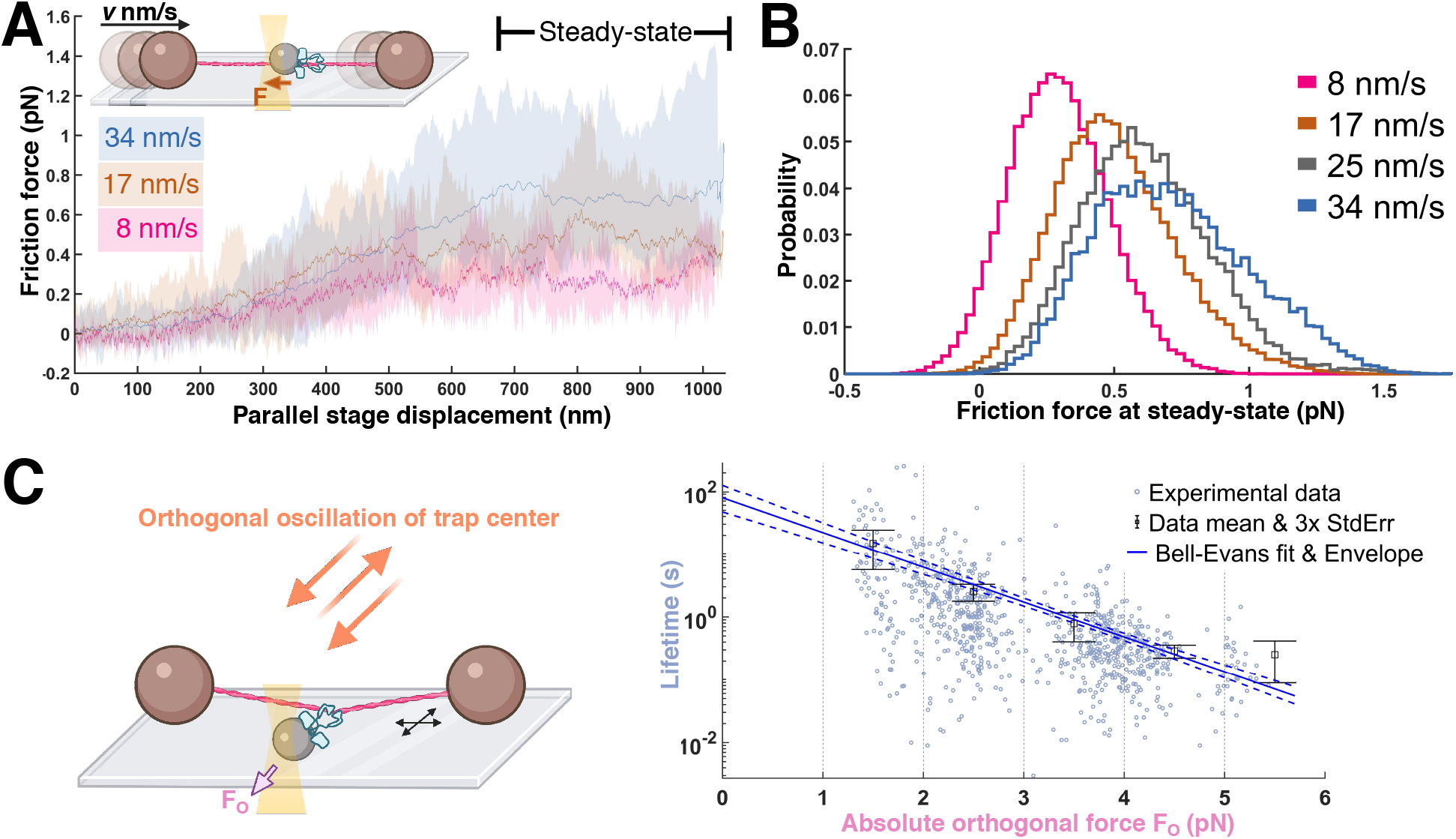
Characterization of the minimal sliding complex. **(a)** Constant stage velocity experiment. *Inset*: The stage is moved at a triangular wave pattern at a set speed. *Example data*: Average forces for a given phase of a stage displacement for a representative sliding complex (average across 4, 11, 14 ramping periods at 8, 17 and 34 nm/s respectively). Envelopes are bounded by maximum and minimum values. Data were mean-filtered (200 ms window size) prior to processing for this panel. **(b)** Pooled steady-state friction force distribution (9 beads from 9 flow cells; boxcar averaged at 100 Hz; see Methods) for the minimal sliding complex. **(c)** Lifetimes under perpendicular loading of the minimal sliding complex (870 events, from 5 beads from 5 flow cells), with mean and error bars (3 standard errors of mean) for each 1 pN interval. The data are fit by a Bell-Evans slip bond model (Methods) where the unbinding rate constant *r* is force-dependent as follows 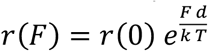 (*F*, force; *T*, temperature; *k*, Boltzmann constant; *d*, distance parameter) with a predicted mean lifetime at zero load of 79 ± 21 s and *d* of 5.2 ± 0.3 nm (errors are standard deviations). The 95% confidence envelope for the fit (blue dashed lines) and standard deviation for each fit parameter were generated through resampling (Methods).

It is likely that ensembles of ezrin-T567D molecules larger than the minimal sliding complex work together *in vivo*. As the bead labeling ratio was increased, we additionally observed sliding assemblies that differed from minimal sliding complexes. These non-minimal sliding assemblies exhibited ∼2-fold or greater friction forces (Fig. 4b; Methods; Supp. Fig. 2), increased heterogeneity in their friction force for a given bead (Supp Fig. 3), and were able to bear ∼4 pN forces in the orthogonal direction for multiple seconds without detaching (Fig. 4a). During constant stage velocity experiments, these “non-minimal” assemblies sometimes exhibited partial, step unbinding and rebinding events, after which they continued sliding (Supp. Figs. 3, 4). Thus, a larger collection of ezrin-T567D molecules than in the minimal sliding complex yielded a sliding connection to F-actin that persisted for multiple minutes and was stable to substantial perpendicular loads.

**Figure 4.**
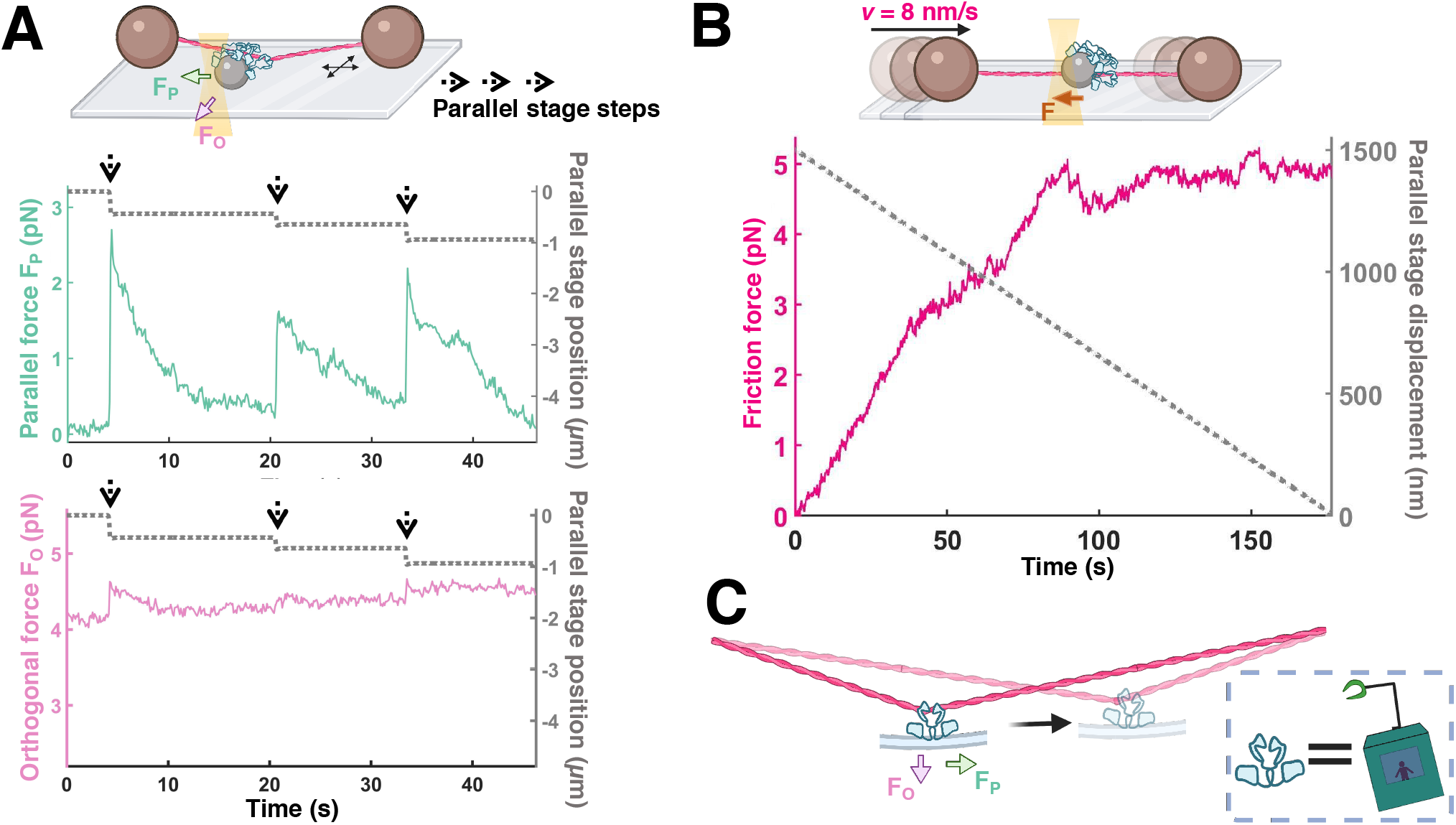
Stronger effective sliding anchors form when ezrin-T567D molecules work together. **(a)** Parallel force relaxation (*green*) by a non-minimal sliding complex under repeated step loading events (*gray*) in the presence of a large, constant orthogonal load (∼4 pN; *magenta*). **(b)** Parallel friction force (stage velocity = 8 nm/s) for a non-minimal sliding complex at steady-state is an order of magnitude larger than that of the minimal sliding complex at the same velocity. All force traces are at 10 Hz. **(c)** Ezrin forms a minimal sliding complex that slides along an actin filament while resisting detachment, similar to a cable car ziplining on a wire.

The ability of the minimal sliding complex to traverse micron distances indicates that it does not follow the F-actin helical pitch, which would require the optically trapped bead to spiral around the actin filament tens of times over this distance. Perhaps relatedly, we noted that the relaxation following a step in parallel force was not smooth, but exhibited bursts corresponding to bead movements of 10-30 nm (Supp. Fig. 5, 6; Methods).

These bursts and pauses potentially reflect the switching of ezrin-T567D contacts from one face of the actin filament to the other, which if present would allow the minimal sliding complex to remain continuously attached while sliding over micron distances.

## DISCUSSION

Previous studies suggest that ERMs play a crucial role in mechanically integrating the actin cytoskeleton to the cell membrane while simultaneously allowing the rapid remodeling of both (*1, 2, 4*) (Fig. 1a). Here we find that a minimal complex comprised of multiple ezrins can slide continuously along actin filaments for tens of seconds and multiple microns while sustaining physiologically relevant perpendicular loads (Fig. 4c). This functionality explains how ezrin can mediate stable attachment of the cell membrane and cytoskeleton while allowing the two to slide relative to each other, for example during bleb retraction (*2, 42*) and compaction in the mouse blastocyst (*31*) (Fig. 1a). Ezrin’s ability to act as a sliding anchor is likewise consistent with its prominent role in microvilli, where it provides stable membrane attachment despite constant treadmilling movement of F-actin toward the microvillus base (*10*) (Fig. 1a).

While we do not know the exact composition of the minimal sliding complex, based on the statistics of our bead labeling, we speculate that it may consist of two ezrin molecules. Future studies are needed to determine the conformation of the minimal sliding complex, and in particular whether it shares the antiparallel organization proposed for non-activated ezrin dimers in solution (*43*). Consistent with our results, quantitative FRAP experiments suggest that *in vivo*, there is an additional kinetic step beyond T567 phosphorylation prior to full ezrin activation (*44*). The implied requirement for recruitment of multiple ezrin molecules suggests that the strength of membrane to cortex attachment may have a nonlinear dependence on the localized membrane density of activated ezrin molecules, a factor that may contribute to ezrin’s prominent roles in sculpting complex membrane geometries (*40*).

To our knowledge, our study constitutes the first systematic examination of the response of an F-actin binding protein to loading at varying angles relative to the actin filament axis. Given these results, it seems plausible that other proteins that slide along polynucleotide and cytoskeletal tracks may exhibit unexpected, angle-dependent responses to mechanical loads (*45*–*52*). In particular, whether other cortical anchors may exhibit variations on the sliding anchoring behavior we characterize here represents an attractive target for future work. It will likewise be interesting to investigate the molecular mechanisms by which ezrin and analogous proteins (*3, 4, 53*) interact with force-generating membrane anchors, for example myosin I, to generate the diverse repertoire of cell shapes and dynamics that characterize animal cells (*1, 2, 54*–*58*). More broadly, our and other studies (*54, 56, 59*) suggest that the emergent properties of the cell cortex may reflect mechanical anisotropies built-in at the level of its individual molecular components, only some of which are currently known.

## ACKNOWLEDGEMENTS

We are grateful to Ms. Amy Wang, Dr. William Weis, Dr. Zev Bryant, and members of the Dunn lab for discussion and comments on the manuscript. E.K. is supported by the Stanford Bio-X Graduate Fellowship and the Biophysics Ph.D. Program at Stanford University. A.R.D. acknowledges the HHMI (Faculty Scholar Award), and the NIH (R35GM130332).

## Notes

### Competing Interest Statement

The authors have declared no competing interest.

