## Supplemental Materials for "The membrane-actin linker ezrin acts as a sliding anchor"

**This PDF file includes:**

Materials and Methods

Figs. S1 to S6

Table S1

References (*34*, *60–65*)

### MATERIALS AND METHODS

Protein expression construct

The following in-frame fusion protein construct was expressed using a pET28a vector: MGSS-6xHis-8xGS-HaloTag-2xG-TEVsite-GGGSGGGGSGGGGSGG-ezrinT567D where TEVsite is the TEV recognition and cleavage site DYDIPTTENLYFQG. The human ezrinT567D sequence used was
MPKPINVRVTTMDAELEFAIQPNTTGKQLFDQVVKTIGLREVWYFGLHYVDNKGFPTW
LKLDKKVSAQEVRKENPLQFKFRAKFYPEDVAEELIQDITQKLFFLQVKEGILSDEIYCP
PETAVLLGSYAVQAKFGDYNKEVHKSGYLSSERLIPQRVMDQHKLTRDQWEDRIQVW
HAEHRGMLKDNAMLEYLKIAQDLEMYGINYFEIKNKKGTDLWLGVDALGLNIYEKDDKL
TPKIGFPWSEIRNISFNDKKFVIKPIDKKAPDFVFYAPRLRINKRILQLCMGNHELYMRR
RKPDTIEVQQMKAQAREEKHQKQLERQQLETEKKRRETVEREKEQMMREKEELMLR
LQDYEEKTKKAERELSEQIQRALQLEEERKRAQEEAERLEADRMAALRAKEELERQAV
DQIKSQEQLAAELAEYTAKIALLEEARRRKEDEVEEWQHRAKEAQDDLVKTKEELHLV
MTAPPPPPPPVYEPVSYHVQESLQDEGAEPTGYSAELSSEGIRDDRNEEKRITEAEKN
ERVQRQLLTLSSELSQARDENKRTHNDIIHNENMRQGRDKYKDLRQIRQGNTKQRIDE
FEAL. Cloning and sequence verification were done by Epoch Life Science.

Protein expression and purification

BL21(DE3) competent bacterial cells were transformed with the ezrin-T567D expression plasmid through heat shock at 42 °C, grown in LB broth and plated on LB agar plates with 50 µg/ml kanamycin to produce colonies. A starter culture of ~5 ml LB broth with 50 µg/ml kanamycin was induced with a colony from the plate, and grown overnight at 37 °C in a shaker at 220-240 rpm. The starter culture was then used to induce a larger volume of LB broth with 50 µg/ml kanamycin, and grown at 37 °C and 220-240 rpm. A small volume was used to make a bacterial stock in 25% glycerol and stored at -80 °C. After reaching an optical density (OD) of 0.8, the culture was induced with 0.10-0.15 mM IPTG for protein expression and grown in at 18 °C in a shaker at 220-240 rpm for ~16 hours.

Cells from the culture were then collected by centrifugation at 6,000 x G for 20 min at 4 °C. The centrifuge bottles were moved to ice and the supernatant media discarded. While on ice, the pellets were resuspended by adding, per pellet of 500 ml culture, a total 10 ml lysis buffer (300 mM NaCl, 50 mM $NaH_2PO_4$, 10 mM imidazole, brought to pH 8 using NaOH) with 14 µg/ml PMSF (50-103-5662, Fisher), 0.2 mg/ml Lysozyme type VI (76177-422, VWR), 7 milli-units/µl DNase I (4536282001, MilliporeSigma), 2 cOmplete™ EDTA-free Protease Inhibitor Cocktail tablets (11873580001, MilliporeSigma).

The suspension was moved to a 360 rotary shaker for 30-45 min at 4 °C, after which a 130 Watt ultrasonic processor (VCX 130, Sonics) was used to further lyse the cells on ice in a 4 °C cold room. Per cycle of sonication, we used a 30 percent amplitude, pulsed as 1 second off and 1 second on for 30 total seconds of sonication. ~10 cycles were done in total and in between cycles, we made sure to wait at least 2 min with occasional swirling to prevent temperature rises, and ensured the temperature remained below ~10 °C using an IR thermometer.

Following sonication, the lysate was centrifuged at 12000 x G for 30 min at 4 °C in a 50 ml Falcon tube. The supernatant was collected, kept on ice, sterile filtered and then incubated with 2 ml HisPur™ Ni-NTA resin (88221, ThermoFisher) slurry per supernatant of 500 ml culture ~1.30-2 hours at 4 °C on a 360 rotary shaker.

A gravity flow column with 5 ml capacity (29922, ThermoFisher) was assembled and the bead suspension loaded onto the column, allowing the solution to run through without letting the beads dry. The column was at room temperature while the buffers and elution collection tubes were kept on ice. We ran a total of 3 column volumes of wash A buffer (300 mM NaCl, 50 mM $NaH_2PO_4$, 20 mM imidazole, pH 7.4 with 0.7 mM freshly added β-mercaptoethanol) in 2 continuous rounds of flow, where each round was started when the remaining solution in the column was the minimum amount that still ensured the beads were not dry. This was followed by elution with a total of 8-9 ml using elution buffer (300 mM NaCl, 50 mM $NaH_2PO_4$, 250 mM imidazole, pH 7.4 with 0.7 mM freshly added β-mercaptoethanol), where we collected 0.5-1.5 ml fractions 2 rounds of flow. The fractions enriched in the desired construct, as analyzed in SDS-page, were kept at 4 °C overnight, brought to 1 mM DTT, pooled and loaded on a Superdex 200 Increase 10/300 GL column (GE Healthcare) for Size Exclusion Chromatography (SEC), which was run using the protein storage buffer (20 mM Tris, 150 mM NaCl, pH 8, sterile filtered, freshly brought to 1 mM DTT). Fractions from the saved peak showed the highest UV 280 absorbance and a UV 260/280 absorbance ratio of ~0.6, indicative of minimal, if any, DNA contamination. SDS-PAGE analysis of different fractions showed that the saved peak was maximally enriched in the desired protein construct. Saved fractions were aliquoted and snap frozen to be used in initial experiments.

A second protein preparation (used in analyzed experiments) was done as above, but with the following modifications: Induction with IPTG was done at an OD of 0.6-0.7 after cooling down the culture at 4 °C for 5 min in between moving the culture from 37 °C to 18 °C, where it was then kept on a shaker for ~16 hours. Following protein expression, the centrifuged cell pellets were kept at -80 °C for a week. The lysis buffer was slightly modified in that the DNase I concentration was increased to 10 milli-units/μl and one cOmplete™ EDTA-free Protease Inhibitor Cocktail tablet was used per pellet of 500 ml culture. The Ni-NTA affinity purification step was done in a 4 °C cold room, where the wash step was modified to include 3 wash steps: 7 ml wash A buffer, 7 ml Wash B buffer (Phosphate-Buffered Saline, 1 M NaCl, 0.002% to 0.005% Tween, sterile filtered) and finally 7 ml wash A buffer. In preparation for Ion Exchange Chromatography (IEC), the eluate from the Ni-NTA resin was pooled and buffer exchanged using IEC buffer 1 (20 mM Tris, pH 8, sterile filtered) through two spin and dilution steps at 4 °C with Amicon Ultra-2 ml 10 kDa centrifugal filters (MilliporeSigma), which we estimate to have resulted in a ~3 fold dilution in the salt concentration. The protein solution was sterile filtered and loaded on an anion exchange column (Mono Q™ 5/50 GL, GE Healthcare) in IEC buffer 1, and subsequently eluted with a linear gradient of IEC buffer 2 (20 mM Tris, 0.98 M NaCl, pH 8, sterile filtered). Following SDS-PAGE analysis, two consecutive 300 μl fractions eluting at ~220 mM NaCl were found to be enriched for the desired construct, with minimal contaminants of differing molecular weights. These fractions were pooled and loaded onto a Superdex 200 Increase 10/300 GL column (GE Healthcare) equilibrated with protein storage buffer for SEC. Five consecutive 200 μl fractions corresponding to protein of the anticipated molecular weight were pooled and saved. The saved protein solution (~600 nM) was stored at 4 °C overnight and aliquoted and snap frozen into -80 °C the following day, that is two days after Ni-NTA affinity purification step.

Proteins from both purifications exhibited binding and sliding behavior that was qualitatively similar (data not shown). Tris buffers used in protein purification were brought to the desired pH either by mixing equimolar solutions of tris base (BP154-1, Fisher) with tris hydrochloride (BP153-1, Fisher) or through titration with hydrochloric acid. IEC and SEC were conducted using a GE Akta PURE Fast Protein Liquid Chromatography system at the Stanford ChEM-H Macromolecular Structure Knowledge Center.

Optical trap setup

We performed our optical trap experiments on a commercial Lumicks C-Trap, which uses a 10 W infrared (1064 nm) laser focused through a Nikon 60x objective (CFI Plan Apo, NA 1.2) to produce two traps with one (Trap 1) more sensitive than the other (Trap

2). An epifluorescence imaging setup was added to image fluorescent actin filaments using a 532 nm laser (Coherent OBIS 532-80-LS) and a sCMOS camera.

Buffer components

The sources of following chemicals were Fisher: $MgCl_2$ (600-30-96), $CaCl_2$ (C79-500), KCl (P217-500), Tris base (BP154-1). Tris buffer stocks were brought to the desired pH through titration with hydrochloric acid (except for protein purification; see above). F-buffer was prepared as 20 mM Tris, 50 mM KCl, 2 mM $MgCl_2$, 0.2 mM $CaCl_2$, pH 8, sterile filtered and either kept at 4 °C or stored aliquoted at -80 °C. 10x F-buffer was prepared as 10 times more concentrated in all components of F-buffer, sterile filtered and stored aliquoted at -80 °C. FBSA buffer was prepared as F-buffer brought to 1 mg/ml ultrapure Bovine Serum Albumin (BSA; MCLAB, UBSA-100) and either directly used while storing at 4 °C for ~2 days or stored aliquoted in -80 °C. DTT (DTT100, GoldBio) was dissolved in water, sterile filtered and stored as 1 M aliquots at -80 °C. Two sources of ATP were both stored as aliquots at 100 mM at -80 °C. ATP from Calbiochem (1191) was dissolved and adjusted to pH 8 with NaOH. ATP from ThermoFisher (R0441) was bought in solution form already adjusted to pH 7.3-7.5 with NaOH and stored at -20 °C before aliquoting into -80 °C.

PI(4,5)$P_2$ diC4 (P-4504, Echelon Biosciences) was either dissolved directly in F-buffer and stored at -80 °C (as aliquots or as stock), or dissolved in ultrapure water, stored at -80 °C and diluted 20x in F-buffer before usage (see Tightrope assay for details of usage). Phalloidin was stored in aliquots at -80 °C dissolved to 1 mM in water or F-buffer. Two sources of phalloidin were used, at least one of which was Cayman Chemical (NC1108931, FisherScientific).

Trolox (648471, MilliporeSigma) was dissolved in F-buffer to 120 mM and stored aliquoted at -80 °C. For oxygen scavenging, we used the pyranose oxidase and catalase system (*60*) (POC). Glucose (anhydrous dextrose; BP350500, Fisher) for use with POC was dissolved to 60% in F-buffer, sterile filtered, and stored aliquoted at -80 °C. For initial experiments we prepared a stock solution of 750 units/ml pyranose oxidase (P4234-250UN, MilliporeSigma), 100 kU/ml catalase (C40-100mg, MilliporeSigma) and 5 mg/ml BSA (UBSA-100, MCLAB) in 20 mM tris, pH 8 which was sterile filtered, aliquoted, snap frozen and kept at -80 °C. These aliquots were either directly used to add POC in forming the trapping buffer or diluted 2x in F-buffer or FBSA before the addition (see Tightrope assay). For experiments resulting in data presented in this paper, the stock solution concentrations of pyranose oxidase and catalase were halved, BSA was included at ~0.15 to 0.5 mg/ml, and F-buffer was used as the solvent. The stock solution was sterile filtered, aliquoted, snap frozen and kept at -80 °C.

Preparation of fluorescent biotinylated F-actin

Lyophilized rhodamine phalloidin (PHDR1, Cytoskeleton) was resuspended to ~800 µM using 8.7 µl methanol (ACS Spectrophotometric Grade, ≥99.9%, Honeywell Riedel-de

Haën™), rapidly aliquoted in ~0.5 µl volumes into tubes and stored in -20 °C, to be later mixed with F-actin as below.

Actin was purified from rabbit skeletal muscle, stored and biotinylated using biotin-NHS (203118, Sigma) exactly as previously described (*59*). The biotinylated actin was snap frozen at a concentration of ~1 mg/ml (24 µM) in ~20 µl aliquots in G-buffer (5 mM Tris pH 8.0, 0.2 mM $CaCl_2$, and 0.2 mM ATP) with 1 mM DTT. Before polymerizing biotinylated actin, an aliquot was thawed on ice for ~30 min, and 20 µl from the aliquot was centrifuged in a TLA100.2 rotor at 60k RPM for 10 min at 4 °C to remove aggregates. The supernatant was moved to a plastic tube, during which the total remaining volume was estimated. 10x F-buffer containing 10 mM DTT and 10 mM ATP was then added to the tube at a volume 1/9$^{th}$ that of the supernatant, inducing polymerization at the actin concentration of ~22 µM. This was mixed and polymerized while on a rotator at room temperature for ~40 min, after which it was diluted to 110 µl (~3.5 µM) using F-buffer with 1 mM DTT and 1 mM ATP, and transferred to a tube of a rhodamine phalloidin aliquot containing 0.5 µl of ~800 µM rhodamine phalloidin. This fluorescent biotinylated F-actin stock was kept on ice at 4 °C for 1-2 days for rhodamine phalloidin to incorporate into filaments, after which it was kept on ice at 4 °C and used in the tightrope optical trap assay within ~2-3 weeks.

<u>Tightrope assay</u>

*Functionalization of trapping beads*

Unless stated otherwise, the beads prepared with the below procedure were used in optical trapping. All centrifugations were done at 3000 x G for 5 min on a benchtop centrifuge, and all sonication steps were performed with a bath sonicator. When removing supernatants from bead pellets, care was taken to leave a minimal amount of solution remaining in order to keep the beads wet.

BSA (UBSA-100, MCLAB) was functionalized with Halo-ligand (HaloTag® Succinimidyl Ester (O4) Ligand; P6751, Promega). Fresh Halo-ligand was thawed to room temperature before opening its vial and dissolved to 80 mM in anhydrous DMSO (dimethyl sulfoxide; 900645, MilliporeSigma) from a freshly opened ampule. This was then immediately mixed with a 100 µM BSA solution in phosphate-buffered saline (PBS, pH 7.4) to achieve 3 mM Halo-ligand at less than 4% DMSO per reaction tube. The experimental reactions were paired with control reactions in parallel where the DMSO contained no Halo-ligand. The reaction mixture tubes were incubated for 2.30 hours at room temperature on a high-angle shaker, and 3.30 hours at 4 °C on a 360 rotator. During the incubation at 4 °C, samples from each of the experimental and control reaction mixtures were desalted (PD Minitrap™ G-25; GE28-9180-07, MilliporeSigma) and further reacted with a HaloTag fused protein which showed >1 new molecular species other than BSA and the HaloTag fused protein in SDS-PAGE analysis, indicative of multiple Halo-ligand links per BSA molecule. Consistently, a high labeling efficiency of BSA with Halo-ligand was suggested by the noticeable shift in apparent molecular weight as analyzed through SDS-PAGE. Aliquots of the reaction mixture were snap frozen and stored at -80 °C, where the non-desalted aliquots were used in the reactions with beads as described below, referred to as BSA-Halo-ligand for the experimental and BSA-Control for the control solution aliquots. The desalted aliquots were used in SDS-PAGE analysis to recheck the high labeling efficiency of BSA, to confirm the preservation of its cross-linking activity to HaloTag fused proteins, and to confirm the functionality of the HaloTag in the ezrin construct.

BSA-Halo-ligand was further functionalized onto beads to be used for optical trapping. First, we activated carboxyl silica beads (mean diameter 1.0 µm; SC04000, Bangs Laboratories) with EDC (1-ethyl-3-(3-dimethylaminopropyl)carbodiimide hydrochloride; PG82079, ThermoFisher) and Sulfo-NHS (N-hydroxysulfosuccinimide; PG82071, ThermoFisher) as follows: Carboxyl silica beads were resuspended at 30 mg/ml in MES buffer (sterile filtered 0.1M MES, 0.9% sodium chloride, pH 4.7 made in ultrapure water with BupH™ MES Buffered Saline Packs; 28390, ThermoFisher), vortexed and sonicated for 15 min. This batch was then split and diluted to 9 mg/ml in 1 ml MES buffer per tube. The following wash procedure was done 3 times per tube: 30 second sonication, centrifugation to pellet the beads, removal of supernatant, and resuspension to 9 mg/ml in MES buffer. After an additional 30 second sonication, first Sulfo-NHS and then EDC, each freshly and separately dissolved in MES buffer at 190 mM and 230 mM concentrations respectively, were sequentially added to the tubes which had a final concentration of 43 mM Sulfo-NHS, 30 mM EDC and 5.8 mg/ml beads in a final volume of 1.55 ml per tube. Care was taken to open the Sulfo-NHS and EDC containers after fully thawing to room temperature. Each tube was then vortexed, bath sonicated for 2 min, and kept on a high-angle shaker for 15-20 min where additional manual mixing of tubes via inversion and vortexing were done during the incubation, with a 30 second bath sonication towards the end. The activated beads were then centrifuged and resuspended in PBS (pH 7.4) after removal of the supernatant. This was repeated once more, after which beads from all tubes were pooled together, sonicated for 2 min, mixed and then split into tubes for reaction with BSA-Halo-ligand or BSA-Control. The final reaction mixture per tube contained 9 mg/ml beads and 17 µM BSA-halo-ligand or BSA-Control in PBS (pH ~7.4), which was sonicated, vortexed and kept on a high-angle shaker to react for 3 hours at room temperature. Following centrifugation, the supernatant was removed and bead pellet resuspended in PBS with 20-40 mM glycine (sterile filtered, pH ~7.2) to quench the reaction, sonicated for 30 seconds, vortexed, and incubated while mixing for 35 min where 1 mM DTT was included in the last 10 min. The beads were then washed twice with PBS with 1 mM DTT through centrifugation, and finally resuspended for passivation in 0.5% casein (from C4765, MilliporeSigma; stored at 4 °C), 0.5% BSA, 1 mM DTT in a final 85% PBS and 15% water mixture at 9 mg/ml bead concentration. The suspensions were vortexed, sonicated for 2 min and incubated for 2 hours while mixing, with extra vortexing and sonication in middle of the incubation. The beads were washed twice via centrifugation, and finally resuspended at 9 mg/ml beads in 0.1% BSA, 0.05% casein, 1 mM DTT in PBS. The bead solution was mixed, sonicated for 2 min, snap frozen in 40 µl aliquots and stored at -80 °C.

Following thawing, a second round of passivation was performed as follows. Pluronic F-127 was prepared within 3 days at 5% in F-buffer and sterile filtered. BSA-Halo-ligand beads and BSA-Control beads were thawed and each resuspended at 1.4 mg/ml beads in 1% Pluronic F-127, 0.016% BSA, 0.017% casein in a mixture of 80% PBS and 20% F-buffer. The mixture was sonicated for 40 seconds and mixed on a 360 rotator at room temperature for 1-1.30 hours. The beads were then centrifuged and exchanged into 2% casein, 0.5% BSA and mixed on a 360 rotator at room temperature for 50 min. Finally, the beads were washed with PBS through 2 centrifugations and resuspended in PBS that was brought to 2.6 mg/ml beads, 0.1% BSA, 0.08% casein, mixed, sonicated for 50 seconds, snap frozen in 50 µl aliquots and stored at -80 °C. The double passivated BSA-Halo-ligand beads were used in all experiments unless stated otherwise.

We note that only for Batch VII (see Supp. Table 1; next section), the BSA-Halo-ligand beads were prepared differently, with main differences being non-specific attachment of BSA to silica (non-carboxyl) beads, on-bead functionalization of Halo-ligand to BSA, and passivation of beads solely with BSA. We did not exclude this batch in our tallying as we did not observe any qualitative differences in F-actin binding behavior (i.e. binding lifetimes, sliding behavior), and thus used it, with the other batches, in deducing the percentage of sliding and stepwise detaching complexes for the given bead activity level (next section).

*Labeling trapping beads with HaloTag fusion protein*

Here we describe the general protocol for attaching HaloTag fusion proteins to the BSA-Halo-ligand or BSA-Control beads, with specific details per batch given in Supp. Table 1. In summary, bead batches at different ezrin-T567D labeling ratios were made by combining FBSA (see *Buffer components*) with the components for incubation for 2-75 min at room temperature in final volume at 50-200 µl and the following ranges in final concentrations: 4-150 nM ezrin-T567D, 0.4-2 mg/ml beads, ~1 mM fresh DTT. The beads were generally added into the larger volume in which the HaloTag fused protein was already diluted so as to aid in homogeneous labeling (tip from Bangs Laboratories technical library). Bubbles introduced during the initial mixing were removed, which lead to loss of some solution. At the end of the incubation, the mixture was centrifuged at 3000 x G, 5 min. The bead pellet was washed at room temperature by repeatedly removing the supernatant and carefully flowing in 90 ul FBSA with 1 mM DTT without disturbing the pellet, for a total of 1.5 ml FBSA wash. Care was taken to leave a minimal amount of supernatant at each step to keep the beads wet. Finally, by judging the pellet size, the washed pellet was resuspended in FBSA with 1 mM DTT to a bead concentration of ~0.2 mg/ml. The batch resuspension was bath sonicated up to 2 times for ~15 seconds each. The resuspended beads were then kept at 4 °C for up to ~3 hours, during which they were used for experiments, and/or snap frozen in 4-8 µl aliquots. Multiple experiments showed no difference in binding behavior between frozen vs non-frozen beads.

The BSA-Halo-ligand beads we used were confirmed to be well-passivated compared to the BSA-Control beads by the attempted labeling of BSA-Control beads with HaloTag ezrin-T567D, which tested for non-specifically adsorbed species of HaloTag ezrin-T567D that retained F-actin-binding activity. We compared the activity of BSA-Halo-ligand and BSA-Control beads to each other in paired experiments at two different labeling concentrations and found that non-specific labeling of BSA-Control beads with HaloTag ezrin-T567D was negligible, as desired. It is worth noting, however, that we did not notice any qualitative differences in F-actin binding characteristics (i.e. binding lifetimes, sliding behavior) for beads that were deliberately prepared by nonspecific HaloTag ezrin-T567D adsorption (data not shown).

*Reagent preparation*

The stock of 3 µm diameter streptavidin-coated polystyrene beads (CP01005, Bangs Laboratories) was diluted 1:10 in F-buffer for the final working stock (~1 mg/ml beads), after washing and sonication as follows: Each washing step consisted of centrifugation at 3000 x G for 5 min in a tabletop centrifuge and removal of the supernatant, which was followed by resuspension in fresh solution. The bead stock, kept at 4 °C, was first diluted ~1:10 in ultrapure water by pipetting 50 µl of well-vortexed beads into 500 µl ultrapure water in an Eppendorf tube, and kept at the same dilution whenever resuspended after washes. The bead suspension was then washed and resuspended in ultrapure water twice, sonicated in an ultrasonic bath for 5 min, re-washed and resuspended in ultrapure water, sonicated for 5 min, and finally washed and resuspended in FB twice. The tube was then sonicated for ~5 min in an ultrasonic bath. This working stock was kept on ice at 4 °C for 2-3 weeks for use in the tightrope optical trap assay.

Pluronic F-127 used for flow-cell passivation was dissolved at 5% w/v in F-buffer, kept at 4 °C until bubbles were mostly removed and then sterile filtered, and stored aliquoted at 4 °C. For experiments with bead batches I-VI (Supp. Table 1), 5% casein solution (C4765, MilliporeSigma) was aliquoted, snap frozen and stored at -80 °C, while it was stored at 4 °C without freezing for older experiments.

*Flow cell protocol*

Microscope slides (12-544, Fisherbrand™ Premium Plain Glass Microscope Slides), microscope coverslips (48366-227, VWR) and double-sided tape (Scotch) were used to form a flow cell as described previously (*34*), within the day of the experiment. Briefly, two stripes of tape of length ~30 mm were laid parallel on the long axis of the slide to create a channel in between them of width ~5 mm. A coverslip was then placed on top of the tape, and good contact with tape was ensured by pressing on the coverslip-tape contacts with back of a marker. This produced a flow cell that held a volume of ~10-15 µl solution.

At most several hours before optical trap experiments aliquots of the following solutions were placed on ice or a metal cooling block immersed in ice, and discarded within the indicated number of days: 1 M DTT (1 day), 100 mM ATP (1 day), 5% casein (1 day), FBSA (~1-3 days), 120 mM Trolox (1 day), POC (1-2 days), 60% glucose (1-2 days), Pluronic F-127 (~month), 1 mM phalloidin (~month), F-buffer. Aliquots that were not discarded within the day were kept at 4 °C in between experimental days. PI(4,5)$P_2$

diC4 was either used in aliquoted forms that were discarded within 2 days, or used from stocks thawed for brief durations before being re-frozen (see *Buffer components*). At the start of experiments, 1 M DTT was diluted to make a 100 mM working stock in FBSA. F-buffer was used for diluting casein to below 5% when needed.

The trapping bead suspension of 4 to 8 μl was either aliquoted from the freshly labeled batch kept on ice or taken from the snap frozen aliquots ~5 min before start of the first flow cell wash. This aliquot was kept at room temperature, and brought to 13 μM $PI(4,5)P_2$ diC4 by careful mixing with $1/4^{th}$ its volume of 0.05 mg/ml $PI(4,5)P_2$. 4 μl of this mixture was later combined with other components of the flow cell (see below) to yield a final $PI(4,5)P_2$ diC4 concentration of 2.1 μM during optical trapping. The optical trapping of beads was performed in T-buffer, the buffer in the enclosed flow cell (0.84% glucose, 0.8-0.9 mM Trolox, 10 μM phalloidin, 1 mM ATP, 1 mM DTT, 7.50 units/ml pyranose oxidase, 1 kU/ml catalase in FBSA), which was sequentially formed as described below.

First, the aliquots were used to prepare W-buffer, which was 1.19x the concentration of T-buffer, where pyranose oxidase and catalase (see *Buffer components*) was usually added immediately before or during each flow cell preparation. W-buffer without POC was kept on ice and used for 1-3 flow cells. At the last passivation step in flow cell preparation, the W-buffer aliquot for the flow cell was diluted to the working, 1x concentration by the addition of ~4 μl of bead suspension to the ~21 μl W-buffer aliquot. This formed T-buffer with beads.

S-buffer (10 μM phalloidin, 1 mM ATP, 1 mM DTT in FBSA) was prepared in amounts to be used for 1-3 flow cells. Usually during flow cell preparation at around step 3, fluorescent, biotinylated F-actin from the ~3.5 μM stock was diluted in an S-buffer aliquot, which was then added to the flow cell at step 5 below. The concentration of F-actin in this solution is estimated to be ~30-150 nM, which was optimized for each batch of fluorescent, biotinylated F-actin and sometimes for each few days of usage, with the ideal flow cell including dumbbells with long filaments (~5-20 μm) every few fields of view, with minimal extra F-actin.

Before preparing the flow cell for optical trapping, each solution to be added was aliquoted in the amounts to be used and moved to room temperature ~5 min before being flowed in to minimize bubble formation due to temperature changes since the optical trapping experiments were performed at room temperature.

The addition of solutions to the flow cell were as follows:

1. >50 μl F-buffer was added to wash the flow cell
2. 20 μl from the well-resuspended working stock of streptavidin bead solution was added and incubated for 8-12 min for non-specific attachment to the surface.
3. The flow cell was washed and passivated (see below)
4. ~42 μl FBSA was added and incubated for ~2 min for further passivation. Beads were mixed into W-buffer to make T-buffer during this incubation step.
5. ~18 μl S-buffer with F-actin was rapidly added by tilting the slide and ensuring smooth flow.

6. ~20 µl T-buffer with beads was rapidly added.
7. Vacuum grease was used to seal ends of the flow cell after which it was placed in the slide holder for optical trapping

The flow cell was incubated with the coverslip facing down during steps 2 and 3 for attachment of beads to the coverslip, by supporting the edges of the slide using empty pipette box lids.

The 3 different wash and passivation protocols used in step 3 above were one of:
a) 40 µl F-buffer wash, 20 µl 5% Pluronic incubation for ~3 min
b) 20 µl F-buffer wash, 20 µl 5% casein incubation for 1.30-2.30 min, 20 µl F-buffer wash, 20 µl 5% Pluronic incubation for 1.30-2.30 min
c) 20 µl F-buffer wash, 20 µl 0.8-5% casein wash, 40 µl F-buffer wash, 20 µl 5% Pluronic incubation for 3 min

The reason for changes in protocol in step 3 for different experiments was an apparent variability in the fragility of actin filaments and the variability in surface passivation. While casein helped passivate against the sticking of trap beads to the coverslip surface, it often also made actin filaments more brittle. One potential explanation for this is the presence of biotin contaminants common in casein stocks.

During optical trapping (see Optical trap setup), a tightrope, aligned in the x-axis (long axis of the flow cell), was found by scanning using brightfield microscopy to monitor streptavidin beads and epifluorescence microscopy to monitor F-actin simultaneously, after which a trapping bead free in solution was captured into Trap 1 (trapping beads stuck to the surface, if present, were not used). Trap 2 was used to clean the surroundings from other trapping beads or to bring beads to Trap 1 to minimize moving Trap 1. Data to be used for fine calibration during post-processing was collected with a trapping bead at or near the Trap 1 position where experimental data was acquired (see Data processing). The fine-calibrated Trap 1 stiffness was 0.020 to 0.032 pN/nm in the x-axis and 0.021 to 0.034 pN/nm in the y-axis.

*Step loading experiments*. To assay binding lifetimes when non-sliding complexes were loaded parallel to the filament, the bead was first brought in contact with a filament, as detected by the displacement in y-force of the trap when pushing against the filament, where the y-axis is orthogonal to the filament. The bead was then kept pressed against the filament at a ~0-0.3 pN orthogonal force and oscillated along the x-axis by moving the trap center in alternating steps in +x and -x, with pauses in between the steps to check for binding above an absolute force threshold upon which the oscillation was stopped until unbinding to below the threshold (Fig. 1c). For experiments in Fig. 1d, the step heights were 0.35, 0.4, 0.5 or 0.6 µm. Each step was completed within ~10 ms. The force threshold was set to 0.25 pN with the coarse calibration during each experiment, which upon fine calibration per collected dataset (see Data processing) was found to be 0.29 pN on average.

*Constant stage speed experiments*. The stage was moved in a triangular wave in the x-axis at mean ramp speeds of 8.4, 17, 25, 34 (<nm/s variation), with each ramp being of sizes varying from 0.75 to 2 μm.

*Binding lifetimes under orthogonal load for sliding complexes*. To assay binding lifetimes, positively identified sliding complexes, for which multiple cycles of steady-state data had been collected, were subjected to load orthogonal to the filament axis, and the trap center was moved in a step oscillation perpendicular to the actin filament axis, with loading step heights 0.9 to 1.6 μm and loading step completion time ~5 ms. Binding events were scored as occurring when the force on the bead exceeded a threshold of 0.5 pN, which upon fine calibration corresponded to 0.55 pN on average. The center position of the trap and the trap oscillation amplitude was determined before data collection by manually moving the stage perpendicular to the filament axis such that peak forces were 1-4 pN. The bead was asymmetrically positioned such that it would approach the filament orthogonally from one side and barely be flush against it at the end of the oscillation.

We expect the effective tether length between the bead and the filament in the orthogonal direction to be ~1 μm. The trap stiffness is 5 to 10 times lower in the *z*-axis than *x* or *y* (estimated by Lumicks); thus, when the bead center is not in the same plane as the filament, we expect a mismatch of up to ~20% between the orthogonal force and total net force on the bead in the force range we assay, due to displacements of the bead in the z-axis (*63*–*65*).

Data processing and analysis

The Matlab software tweezercalib 2.1 (*62*) was used for fine stiffness calibration of the optical trap where the dependence of hydrodynamic friction on frequency and on the bead's proximity to the coverslip surface was taken into account, the position detector was treated as a low-pass filter with one parameter, aliasing was accounted for and the already small crosstalk between *x* and *y* axes was eliminated. Absolute height of the trapped bead from the surface (typically ~1-3 μm from bottom of the bead) was estimated within ~400 nm using a template pre-created in the Lumicks software for a surface streptavidin bead where the trapped bead distance to the surface was known; to determine the difference in height compared to the template height during the experiment, the stage was moved until selected surface streptavidin bead images matched the template. The other inputs were trapping bead radius (0.5 μm), trapping bead density (2 g/cm$^3$), fluid density (1 g/cm$^3$) and fluid kinematic viscosity (10$^6$ μm$^2$/s). The bead height from the surface for fine-calibration data collected for two beads was not noted. Both beads were determined to be non-minimal sliding complexes. For analysis of these two beads, we assumed a bead height of 2 μm (typical in our experiments) for the fine calibration, which we expect to introduce an inaccuracy of at most ~10% in our force measurement, which does not affect our interpretations.

The data from trap experiments assaying binding lifetimes (parallel or orthogonal) was boxcar averaged to 1000 Hz, and binding events were detected as follows, where we consider successful binding when the complex remains bound >15 ms following the completion of loading. First, all possible timepoints where loading of the complex could happen were found by detecting steps in the trap position through the `ischange` function in Matlab. Using this information, we scanned timepoints that corresponded to 15 ms after the completion of a potential loading. When the force along the relevant axis at this timepoint exceeded threshold A (0.75 pN for parallel and 1.1 pN for orthogonal loading), the event was considered a successful binding, and the lifetime was taken as the time interval starting from this point (that is, after 15 ms) until before the force decreased below threshold B (0.225 pN for parallel and 0.275 pN for orthogonal loading). The force for the binding event was taken to be the average over the lifetime.

To analyze the steady-state friction force of sliding complexes, the turning points of the stage during the triangular wave were either manually or automatically detected to extract the timepoints where ramping phases started and ended. However, if sliding complexes unbound/rebound during a ramp, each section of continuous sliding potentially long enough to reach steady-state were manually selected. Likewise, if the experiment was compromised for part of the sliding event, for example by the presence of a nearby diffusing bead, only the uncompromised part was selected.

The timepoint at which steady-state was reached was determined as follows: The optical trap force timeseries were boxcar averaged to 100 Hz, filtered with a moving mean window size of 200 ms and the first timepoint where the force fluctuated to 0.22 pN in the opposing direction to the ramping was taken. An additional 400/$v$ seconds, where $v$ is the stage speed in nm/s, was added to this timepoint to account for the bead rotation (bead radius 500 nm). An additional 15 seconds was further added to further ensure reaching of steady-state. The force traces corresponding to the resulting steady-state time intervals were analyzed as boxcar averaged to 100 Hz (without any moving mean filtering).

*Labeling ratio of bead batches and their associated data*

Here we describe the different batches of trapping beads labeled with HaloTag fusion ezrin-T567D used in our experiments (Supp. Table 1). As explained in the *Flow cell protocol*, we evaluated three different passivation methods at step 3 and found that the casein-based protocols were best at preventing the sticking of beads to the surface or to streptavidin beads, although not perfect. We do not sample stuck beads; thus, excessive sticking of trap beads is potentially problematic when estimating the percentage of active beads for a batch. Control experiments with unlabeled beads (i.e. BSA-Halo-ligand beads with no HaloTag ezrin-T567D during incubation) showed a ~10% ratio of nonspecific sticking of beads to the surface using a casein-based protocol. Per tallied flow cell, we used this control ratio as a guide and noted the ratio of beads stuck to the surface over the course of the experiment to determine until what timepoint, for a given flow cell, statistics could be safely tallied.

When testing the labeling statistics of beads, we made use of filaments that were taut enough that we could ensure, by pushing the bead against the filament at forces of ~0-0.3 pN, that any active complexes on the bead would likely encounter the filament during the parallel trap oscillation. We used 2 types of scans, short (~30 s) and long (~1 min). These durations were determined empirically during optimization. Long scans were able to detect if a bead was in general active, i.e. whether it contained stepping or sliding molecules. However, due to the faster on-rates of sliding complexes compared to stepping complexes, sliding complexes could be easily detected with ~30 s scans alone. Thus, to speed up bead sampling, sometimes the short scan procedure was used to detect if a bead contained a sliding complex or not, where a sliding complex was further confirmed by manual movement of the stage and/or steady-state friction experiments. In some data sets, we started off using long scans, but then switched to short scans during course of the experiment. In these cases, data from flow cells were divided into two sections during data processing: the first containing the initial long, scans and the second containing the short scans. These are referred to as flow cell sections below.

From the casein-based passivation protocols, 26 flow cell sections were taken for tallying, where 22 had a stuck bead ratio of ~10% and 4 were closer to ~25%. From flow cells with passivation protocols not containing casein, 6 flow cell sections were taken for tallying, with 10-30% stuck bead ratios. The results of our tallying are shown in Supp. Table 1, where we note how a given bead batch was prepared and the ratio of beads that had stepping vs. sliding complexes. Beads that showed solely a single binding event were not counted as active. Such single-event beads were seen when HaloTag ezrin-T567D was incubated with non-functionalized BSA-Control beads in control experiments (see *Functionalization of trapping beads*), and thus may reflect HaloTag ezrin-T567D molecules weakly associated with the passivation layer that are ripped from the bead when subjected to load.

As indicated above, bead batches fall into two categories, those with activity ≤ 0.11 (batches I, II and III) and those with activity ~0.35 (batches IV, V, VI, VII). Combining data for batches I, II and III together and taking weighted averages, we find that 90% were inactive, 7.5% showed stepping complexes, and 2.2% had sliding complexes. For percentages, we calculate an original estimate from long scans as 90%, 7.5% and 2.5% in the same order as above; however, sliding complex percentage (2.5%) could be made more precise by incorporating data obtained from short-duration scans, which as noted above were designed to detect sliding complexes, but not stepwise detachments. Of the beads with stepping complexes, we detected solely single-step unbinding for 70%, and a mixture of single- and double-step unbinding for 30%. From the beads exhibiting sliding complexes, 3 were minimal and 1 was non-minimal. Assuming Poisson statistics and a purely monomeric molecule, at an inactivity ratio of 90%, 9% of total beads are expected to contain single molecules, in reasonable accord with the fraction of beads showing solely single-step unbinding behavior.

Combining data from batches IV, V, VI and VII and taking weighted averages, we find that these beads had 66% inactivity, 27% stepping complexes and 4.5% sliding complexes, where the percentages do not sum to 100% due to the same considerations as above. For reference, at 66% inactivity ratio, with same assumptions as above for a purely monomeric molecule, 27% of beads would be expected to contain a single molecule and 6% are expected to contain two molecules.

We interpret above results, where sliding complexes are rarer than complexes showing stepwise release, to indicate that sliding complexes include multiple ezrin-T567D molecules, and that the minimal sliding complex likely is formed by the association of two ezrin-T567D molecules with F-actin.

We note several issues that may present potential experimental biases in our tallying. Firstly, in some fields of view in our flow cells, we noticed beads that were in aggregates of two or more beads. Such aggregates did exist in the control bead batch (i.e. BSA-Halo-ligand beads with no HaloTag ezrin-T567D during incubation) as well, but anecdotally were more noticeable in bead batches with higher labeling (~65% inactivity). Additionally, in some flow cells except for those using the control bead batch, there were trapping beads already attached on actin filaments. This is not surprising due to the minute-scale no-load binding lifetime of sliding complexes, as well as the long lifetimes that could be expected from multimolecular stepping complexes. We did not include such beads in our tallying since their detection involved observer bias. We noticed a negligible number of beads (total of ~3) pre-tethered to actin filaments in the 18 flow cells in the low activity (≤ 11% activity) batches, while this phenomenon was more common in batches with higher activity (≥ 30% activity) where we noticed a few beads tethered to filaments per flow cell for multiple flow cells. Pre-tethered beads were typically found to be non-minimal sliders. Finally, it is possible that long (~1 min) scans may have still missed a small fraction of the beads with active molecules, leading to a slight underestimation of their numbers.

When not interested in the statistics of bead labeling, we considered all assayed beads for analysis, irrespective of whether they were assayed before or after the stuck bead ratio was above a threshold. As mentioned above, no data was collected from stuck beads. For the analysis of minimal sliding complexes, we only included minimal sliding complexes for which data was collected for in batches II, III, and IV since these batches were most completely characterized. In total, minimal vs non-minimal sliding complexes could be assigned for 21/23 beads where we simply did not collect enough information for 2 beads to confidently ascribe their status as minimal or non-minimal. The partial step unbinding and rebinding events seen in some non-minimal sliding complexes are shown in Sup. Figs. 3, 4. Data from batches II and III (~90% inactivity) was also used to calculate the force-dependent lifetime of single molecules when loaded in parallel, where we analyzed data collected from beads that solely produced single-step unbinding events (Fig. 1).

*Bursts and steps exhibited by minimal sliding complexes*

As described in the manuscript, in relaxation traces from step loading of minimal sliding complexes, sometimes bursts (stalls interspersed with sliding) can be seen, which are also apparent in pairwise distance distribution analysis of the some of the traces (Supp. Figs. 5, 6). While we do observe bursts at both low (~2 pN) and high (~4 pN) forces, we expect our burst size estimates and temporal resolution to be worse for lower forces due to the higher effective compliance.

*Slip bond model fitting*

The Bell-Evans model for a slip bond (*61*) predicts an exponential dependence of the unbinding rate constant $r$ on the applied force $F$ as follows

$$r(F) = r(0)\, e^{\frac{F\, d}{k\, T}}$$

where $T$ is temperature, $k$ is Boltzmann constant and $d$ is the distance parameter. This results in the following exponential probability distribution $P(\tau)$ for bond lifetime $\tau$:

$$P(\tau) = r(0) e^{\frac{F\, d}{k\, T} - r(0)\, \tau\, e^{\frac{F\, d}{k\, T}}}$$

. The expression was fit to the minimal sliding complex binding lifetimes under orthogonal force as described in Fig. 3c legend. The 2.5-97.5% confidence intervals were generated through resampling: the dataset was randomly resampled by the dataset size 1000 times, with each resampling fitted to the slip bond model, and the resulting 2.5 and 97.5 percentile lifetimes at each force value taken.

**Supplementary Figure 1**

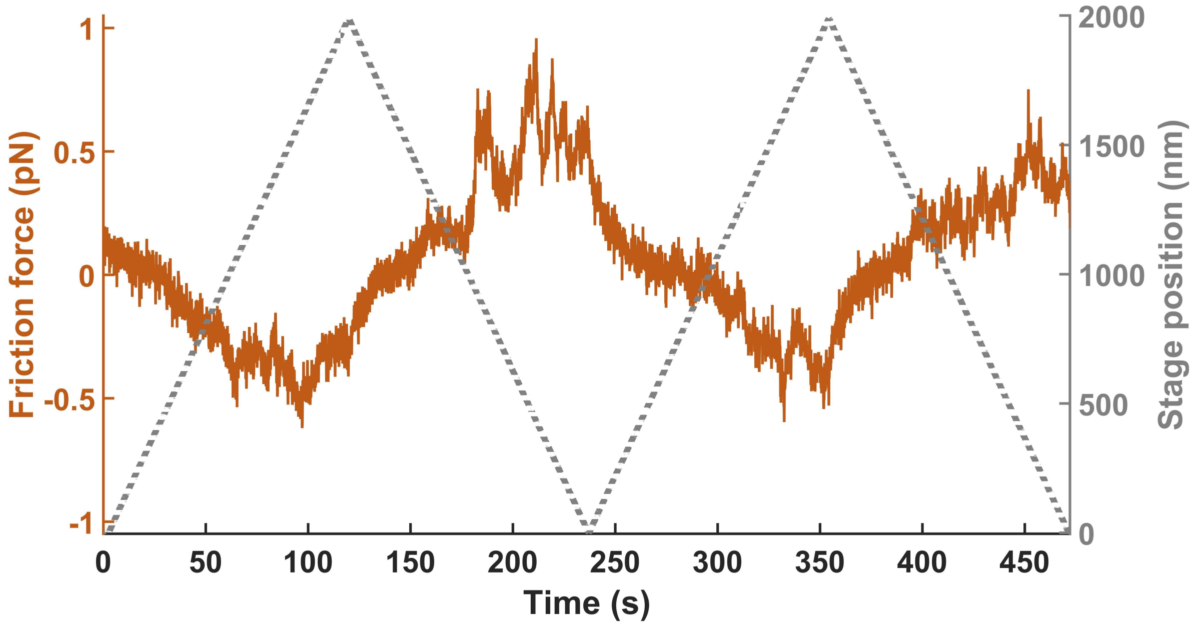


**Supplementary Figure 1. A minimal sliding complex of ezrin-T567D sliding on F-actin for four consecutive stage ramps.** Here, the stage is translated in one direction along the actin filament at 17 nm/s for 2000 nm before reversing its direction to start a new ramp. The minimal sliding complex exerts a friction force in the direction opposite to the stage movement. Force trace is boxcar averaged to 10 Hz.

### Supplementary Figure 2

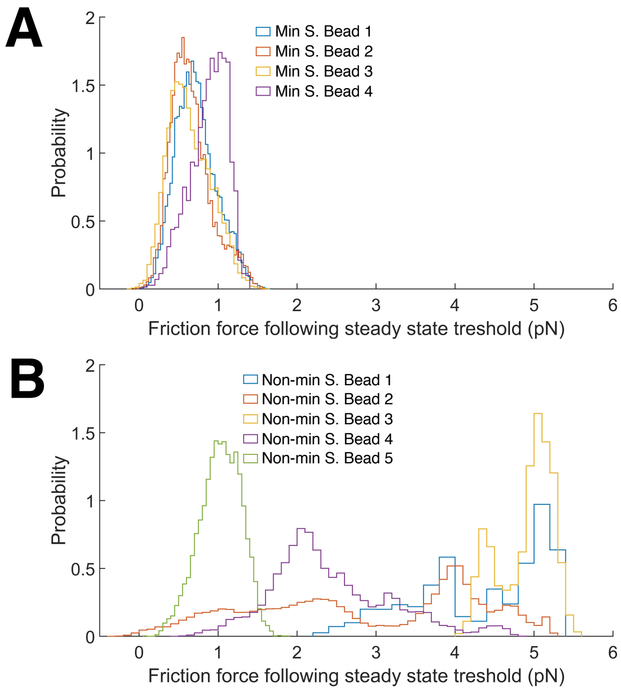


**Supplementary Figure 2. Friction force distributions following steady-state threshold. (a)** The average steady-state friction force distributions for minimal sliding (Min S.) complexes at 34 nm/s after applying steady-state analysis to beads from batch II (Min S. Bead 1), batch III (Min S. Beads 3, 4) and batch IV (Min S. Bead 2) (Supp. Table 1; Methods). **(b)** Non-minimal sliding (Non-min S.) complexes at 34 nm/s exhibit markedly different friction force distributions when the same steady-state analysis is applied (Non-min S. Beads 1, 2, 3, 4 from batch IV; Non-min S. Bead 5 from batch III) due to heterogeneity in their steady-state friction forces (Supp. Fig. 3), and due to taking longer to approach the larger steady-state forces (Fig. 4b; Supp. Fig. 3, 4). While one bead with a non-minimal sliding complex (Non-min S. Bead 5) exhibits a friction force distribution similar to that of minimal sliding complexes at steady-state, it was assigned as non-minimal due to exhibiting multiple cases of partial step unbinding when approaching high forces, which is a phenomenon seen in other non-minimal sliding complexes (Supp. Fig. 4) but not in minimal sliding complexes in constant stage ramp experiments.

**Supplementary Figure 3**

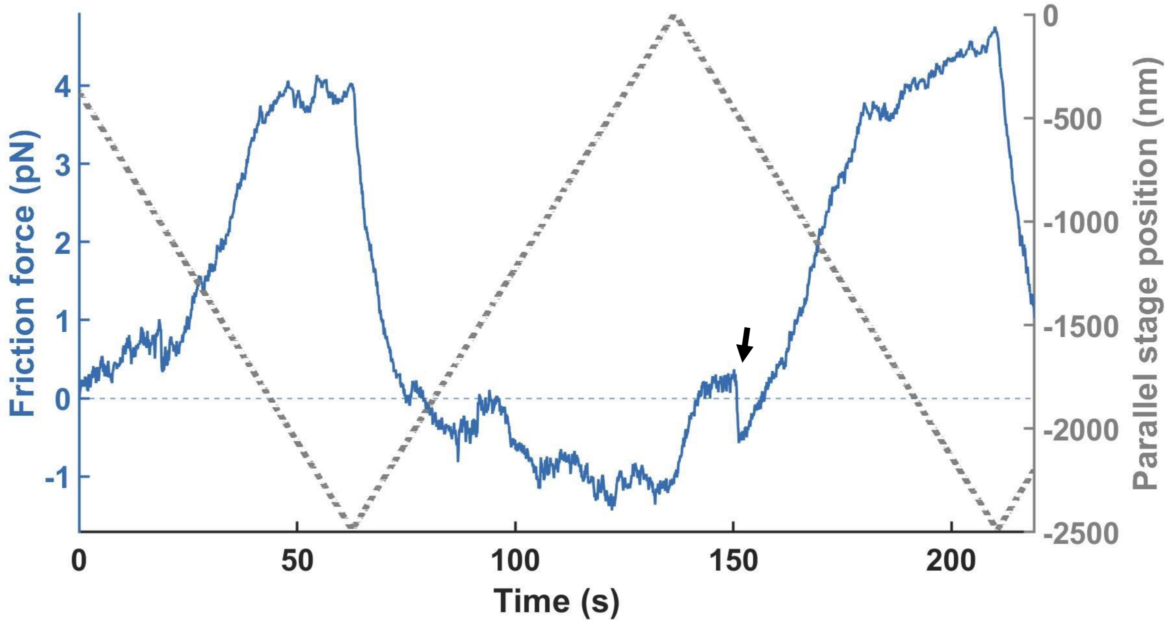


**Supplementary Figure 3. Non-minimal sliding complexes can exhibit larger and heterogeneous friction forces compared to minimal sliding complexes.** A non-minimal sliding complex is seen to approach different steady-state friction forces when the stage is ramped at 34 nm/s in opposite directions along the filament. This may reflect different numbers of ezrin-T567D molecules bound to F-actin when the bead leans in a given direction. The force step indicated by the arrow at ~150 s can be interpreted as the rebinding of an additional complex. This would be consistent with the higher friction force reached in the rest of the ramp (~4 pN at ~190 s compared to ~1 pN at ~130 s in the opposite direction). Force trace is boxcar averaged to 10 Hz.

Supplementary Figure 4

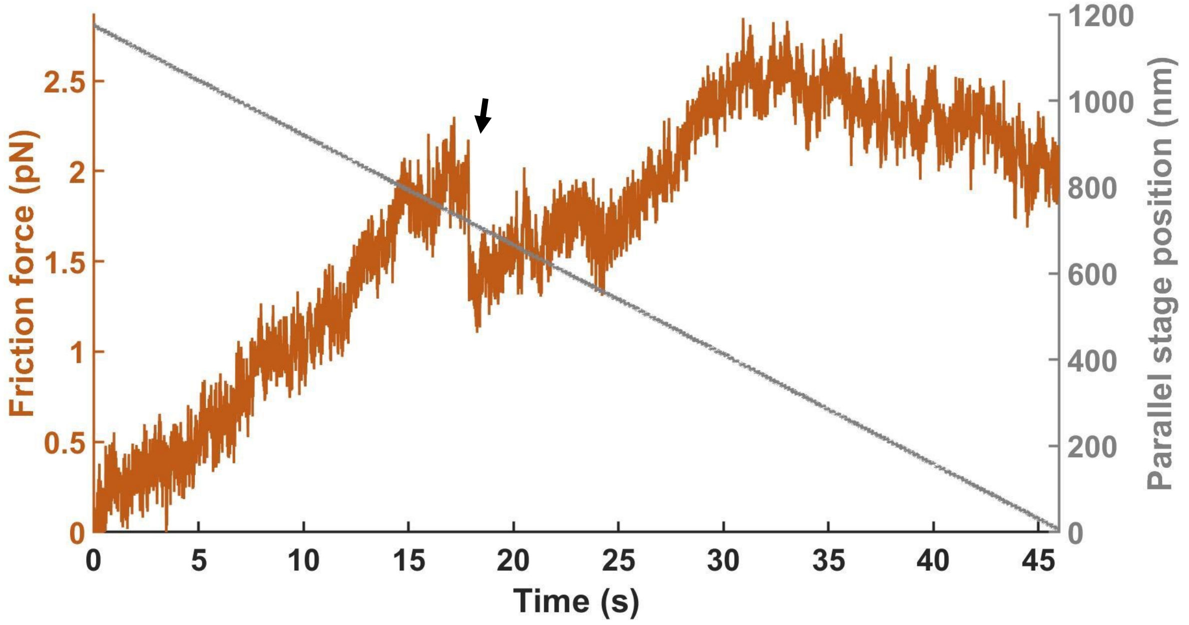


**Supplementary Figure 4. Friction force of a non-minimal sliding complex at a constant stage velocity of 17 nm/s.** An example partial unbinding event of magnitude ~0.5 pN at ~17 s during sliding is indicated with an arrow. Force trace is boxcar averaged to 100 Hz.

**Supplementary Figure 5**

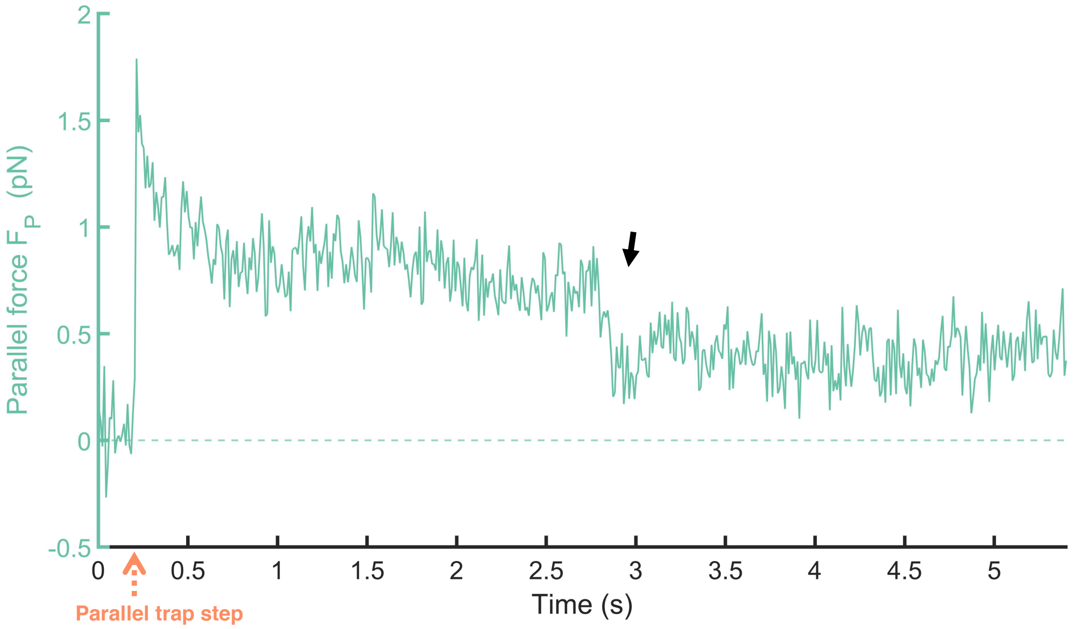


**Supplementary Figure 5. Minimal sliding complexes can exhibit >10 nm bursts during sliding.** A step perturbation parallel to the filament is applied by a step movement of the trap along the filament at ~0.2 s (orange arrow; trap position is not shown) increasing the parallel force to ~1.5 pN, after which the force relaxes though complex sliding. An example ~20 nm burst (arrow). Force trace is boxcar averaged to 100 Hz. Compare to Fig. 2a and Supp. Fig. 6.

**Supplementary Figure 6**

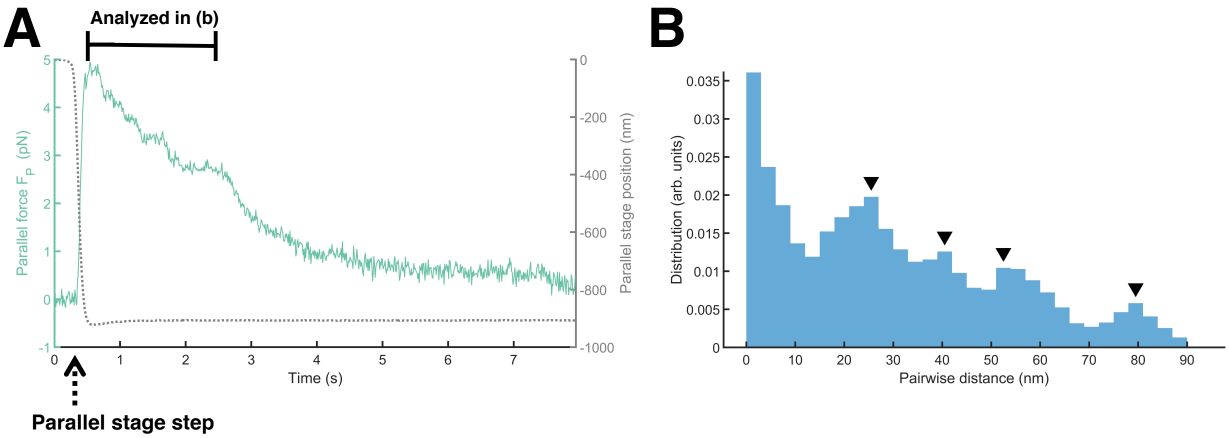


**Supplementary Figure 6. Pairwise distance analysis of >10 nm bursts of a minimal sliding complex during sliding. (a)** A step perturbation raises the parallel force to ~5 pN, after which the force on a minimal sliding complex relaxes to close to zero. **(b)** The pairwise distance distribution corresponding to ~0.5 to 2.6 s in (a). Estimates of burst sizes are indicated by the first peak at 25.5 nm and the following inter-peak distances of 15 nm, 12 nm, 27 nm, as indicated by triangles.

**Supplementary Table 1**

| HaloTag fusion ezrin-T567D concentration at incubation (nM) | Bead concentration during incubation with ezrin-T567D (mg/ml) | Incubation time (min) | Batch ID | Long Scan Active bead fraction | Long Scan Stepwise detaching bead fraction | Long Scan Sliding bead fraction | Long Scan Total assayed beads | Short Scan Sliding bead fraction | Short Scan Total assayed beads |
|---|---|---|---|---|---|---|---|---|---|
| 12 | 2.04 | 2 | **I** | 0.06 | 0.06 | 0.00 | 16 | | |
| 49 | 0.48 | 40 | **II** | 0.10 | 0.10 | 0.00 | 42 | | |
| 26 | 0.5 | 75 | **III** | 0.11 | 0.06 | 0.05 | 62 | 0.02 | 65 |
| 23 | 0.5 | 37 | **IV** | 0.35 | 0.29 | 0.05 | 75 | 0.03 | 267 |
| 131 | 0.41 | 32 | **V** | 0.30 | 0.10 | 0.20 | 10 | | |
| 129 | 0.56 | 36 | **VI** | 0.32 | 0.25 | 0.07 | 28 | | |
| 4 | 1.1 | 15 | **VII** | 0.38 | 0.29 | 0.08 | 24 | | |

**Supplementary Table 1. Preparation details and bead batch activity statistics.** Labeling conditions for different batches of optical trap beads. All BSA-Halo-ligand beads were functionalized with Halo-ligand with the same protocol except for Batch VII where a different protocol was used (Methods). For each bead batch, we collected bead activity statistics by scanning multiple beads to estimate the fraction of active beads, as well as the fraction of beads exhibiting stepwise detachment or sliding. A Long Scanning protocol (LS) determined whether and how a bead was active (i.e. stepwise detachment or sliding) while a short scanning protocol only determined whether a bead exhibited sliding or not (Methods).
